# Preparatory Bias Currents Alone Write Letters and Words in a Fixed Recurrent Circuit

**DOI:** 10.64898/2026.11.13.766302

**Authors:** Alexander James White, Chung-Chuan Lo

## Abstract

We show that preparatory input currents, or bias currents, alone can make a recurrent circuit with fixed weights write all 52 cursive letters, and words built from them. Each letter is produced by its own bias pattern, ramped in as a preparatory drive and withdrawn at a go cue; the recurrent weights and the readout never change, and the readout’s weights are random, with a single global gain picked on four letters (a–d). We take the stability-optimized circuits (SOCs) of Hennequin *et al*. and extend them with mutual inhibition: all inhibitory neurons inhibit one another through a single shared weight (MISOC, mutual-inhibition SOC). With 200 units, the pen of the SOC stays on average 1.7% of a letter width from the target over all 52 letters, and that of the MISOC 0.9% (25 random circuits; *R*^2^ = 0.996 and 0.999). The other 48 letters are learned through their bias patterns alone, without changing a single synapse, so new letters cannot disturb old ones. Letters learned one at a time chain into words in one continuous run. Mutual inhibition improves the circuit: with a shared weight of 0.5 the MISOC writes with 16–48% less error than the SOC at every network size tested (50 to 200 units), and with fifty network-derived input channels instead of one bias per neuron it keeps its advantage. Bias currents and mutual inhibition let small, fixed circuits hold a large and growing motor repertoire and switch within it without any synaptic change.

## I. Introduction

**A**nimals change what they do in fractions of a second, while synaptic plasticity acts over minutes to days. Across motor cortex, movements are generated by the same circuit with the same connections; what differs between movements is how the circuit is driven [1], [2]. Tonic input, held constant while a behavior is prepared, is a natural control channel. Neuromodulators reconfigure small circuits without rewiring them [3]; contextual input selects computations in prefrontal cortex [4]; a context-dependent tonic input reconfigures the dynamics that time movements, whose speed is then set by the initial state [5]; and in our own work, bias currents alone switch a four-neuron circuit between 24 logic functions [6], while mutual inhibition between inhibitory neurons enlarges the set of functions a circuit can reach [7]. We call this *bias-controlled reconfigurability*: the weights stay fixed, and only the bias to each neuron changes. Whether it scales from discrete computations in tiny circuits to continuous, high-dimensional motor output is an open question.

Two lines of work point toward this. Learning only biases is known to be powerful [8], [9], and a movement-specific preparatory state, set by an input and released at movement onset, is a leading account of how motor cortex selects a movement [1], [10]–[12] So far, however, models have had either simple outputs or heavy machinery. Recurrent-network models of preparatory input, and networks with learned biases, produce simple outputs: a few reaches or a handful of one-dimensional traces [8], [11]–[13] The model that writes many letters from a learned input uses a long short-term memory (LSTM) network with tens of thousands of trained weights, trained on hundreds of samples per letter [9]. And in the preparatory-state models the readout is fitted to the movements: Hennequin *et al*. [10] fix the preparatory states and learn the readout weights by least squares, and Stroud *et al*. [11] fit all readout weights by least squares before learning new movements. Here the biases do all of the learning. The readout is a random pattern that is never fitted, and its only parameter, a single global gain on the output weights, is picked on four letters (a–d); the recurrent weights are built without ever seeing a letter, so this gain is the only synaptic parameter that is set with any letter in view. We combine complex output with simple machinery: a generic recurrent circuit with every weight fixed holds a large repertoire of complex, continuous movements selected by a learned bias alone, each state is reached by an input the circuit actually receives, separately learned movements compose into sequences without relearning, and mutual inhibition between inhibitory neurons makes the circuit write better. We return to how this relates to previous models in the Discussion (Section IV-B).

Here a preparatory bias pattern, ramped in during preparation and withdrawn at the go cue, sets the initial state and is the only thing that differs from movement to movement, and we make this route the subject of a full study. Every state here is *reached*: the bias ramp, delivered during one second of preparation, drives the circuit into it, with the spread of the state limited, the pen asked to stay still throughout preparation (output-null [14]) and brought to rest after the letter. With a random readout whose single global gain is picked on four letters, one fixed circuit writes all 52 two-dimensional cursive letters, across 25 random circuits at each of 5 sizes (*N* = 50, 70, 100, 150 and 200 units); letters learned alone chain into words and whole alphabets in one continuous run; the inhibitory structure of the circuit decides how well it writes; and the bias can be compressed to a few dozen input channels. Our contributions are:

1. a fixed stability-optimized circuit and a fixed random readout write all 52 cursive letters from learned bias patterns alone; the readout’s weights are random and its only parameter, a single global gain, is picked on four letters, so 48 letters are written without any weight, recurrent or readout, ever being fitted to them;
2. letters learned one at a time, each in isolation, chain into words and whole alphabets in one continuous run, and new letters never disturb old ones because nothing shared is changed;
3. all-to-all mutual inhibition between inhibitory units through one shared weight (MISOC) improves accuracy;
4. circuits of a few dozen neurons suffice, and the bias can be compressed: with 50 fixed, network-derived input channels instead of one bias per neuron the MISOC keeps its advantage.

## II. Model and Methods

### A. Overview

The idea of the model is that learning a movement should change nothing that the movements share. A recurrent network with fixed weights generates the dynamics, a fixed random readout turns its activity into pen motion, and each letter has its own preparatory bias pattern: a constant input to every unit that is ramped in while the circuit prepares and withdrawn at a go cue. Learning a letter means finding that bias pattern, and nothing else (Fig. 1). The subsections below describe, in turn, the network, the two circuit types we compare, the letters and the readout, how a bias pattern is learned, how letters are combined into words, and the draws and the input-channel test.

**Fig. 1.**
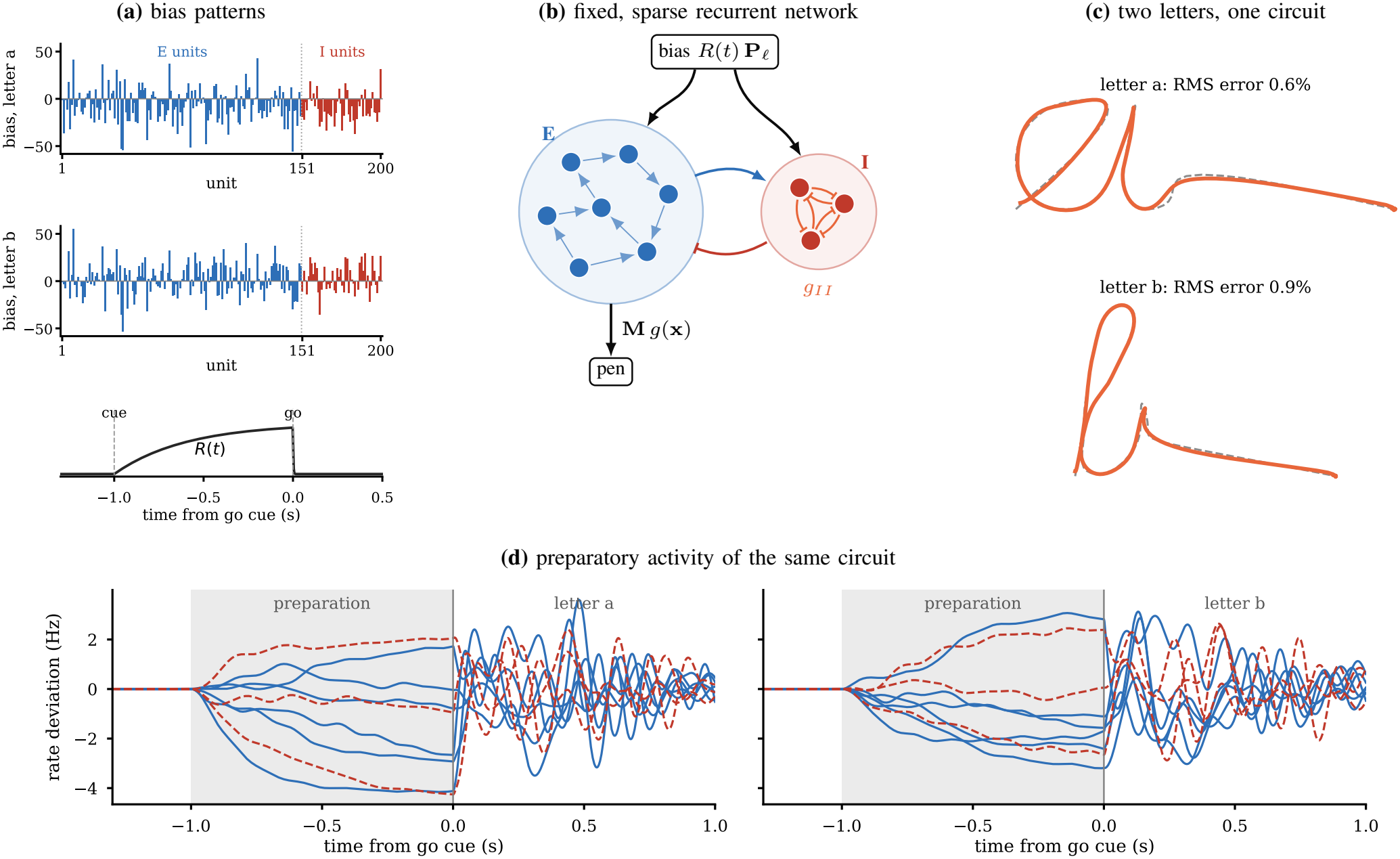
Bias-controlled writing (*N* = 200, MISOC, first draw). (a) Each letter *l* has its own bias pattern **P**_*l*_, one amplitude per unit (blue: excitatory, red: inhibitory units), delivered through a common ramp *R*(*t*) that rises after the cue and is withdrawn at the go cue; shown are the learned patterns of a and b. (b) The recurrent network is fixed and sparse. In the SOC the inhibitory-to-inhibitory weights are optimized synapse by synapse; in the MISOC every inhibitory unit inhibits every other through one shared weight *g*_*II*_ = 0.5. A fixed random readout of the excitatory units moves the pen relative to an arm that carries it one letter width to the right during each letter. (c) The two letters written by this circuit over the letter and the hold (pen: orange; target: dashed). (d) Rate deviations of six excitatory (blue, solid) and three inhibitory (red, dashed) units while the circuit prepares and writes a (left) and b (right): during preparation (shaded) each unit settles at its own preparatory level, set by the bias pattern, and after the go cue the circuit’s own dynamics take over. Weights and readout are identical; only **P**_*l*_ differs.

### B. Network

The circuit is a recurrent network of firing-rate units, driven by a bias pattern that a preparatory ramp *R*(*t*) switches on before the movement and off at the go cue. We use the rate model of Hennequin *et al*. [10]. The potentials **x** of *N* units obey

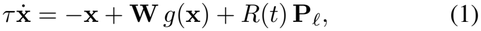

with *τ* = 200 ms, and the firing-rate deviation from a 5 Hz baseline (is *g*(*x*) = *r*_0_ tanh(*x/r*_0_) for *x <* 0 and (*r*_max_ *−r*_0_) tanh *x/*(*r*_max_ *− r*_0_) for *x ≥* 0, with *r*_0_ = 5 Hz and *r*_max_ = 100 Hz. We test *N* = 50, 70, 100, 150 and 200 units, 75% of them excitatory; figures of a single circuit and the numbers quoted for one size use *N* = 200 (150 excitatory and 50 inhibitory units). Simulations are deterministic (no noise) and integrated with forward Euler steps of 0.5 ms.

### C. Stability-optimized circuits

A preparatory state is only useful if the circuit turns it into a rich, movement-length transient, which needs strong recurrence, but strong recurrence in a random network is chaotic. Stability-optimized circuits (SOCs) [10] resolve this: their inhibition is tuned so that the network is strongly amplifying yet stable. To build an SOC, following [10], we start from a sparse random matrix (connection probability 0.1, inhibitory weights three times stronger than excitatory ones on average, spectral radius 10), which is deeply chaotic. Stability is measured by the spectral abscissa *α*(**W**), the largest real part of the eigenvalues of **W**: activity returns to rest only if *α <* 1, and the further below 1, the faster. Because *α* itself cannot be lowered smoothly by gradient descent, we lower its smoothed version [15], a differentiable upper bound on *α*. The excitatory-to-excitatory weights are frozen, and the remaining weights descend this smoothed abscissa while keeping Dale’s law, the mean weight of each block and an inhibitory connection density of at most 40%. Hennequin *et al*. keep all excitatory weights fixed, but at our 75/25 split that leaves the circuit unstable (*α≈* 3.8), so in both circuits we also tune the existing excitatory-to-inhibitory synapses, one by one.

To ask whether the structure of inhibition matters, we build two circuits from the same starting matrix with the same optimization; they differ only in how the inhibitory units connect to one another. In the *SOC* these inhibitory-to-inhibitory weights are optimized synapse by synapse, and the optimization stops when the spectral abscissa *α*(**W**) falls below 0.5. In the *MISOC* (mutual-inhibition SOC) the block is all-to-all mutual inhibition with one shared weight, *W*_*ij*_ = *−g*_*II*_ for *i* ≠ *j*, and only the excitatory-to-inhibitory and inhibitory-to-excitatory weights are optimized. A uniform all-to-all block places the inhibitory difference modes near +*g*_*II*_, so the MISOC cannot reach an abscissa much below *g*_*II*_; we use *g*_*II*_ = 0.5 and optimize for up to 20 000 steps toward *α <* 0.6. At *N* = 200 this gives *α* = 0.745 (SOC: 0.494). The SOC and MISOC of a draw share the same random initial matrix. The higher abscissa does not explain the MISOC’s advantage: SOCs stopped at the MISOC’s abscissa write like fully optimized ones (Section III-D, Supplementary Section S1).

### D. Letters, arm and readout

The task is to write 52 cursive letters, each a two-dimensional pen trajectory of 500 ms. The targets are the letters A–Z and a–z of the Hershey Script Simplex font [16], a single-stroke (centerline) cursive font whose glyphs are pen paths, each smoothed with a cubic spline, resampled to 100 points and shifted to start at (0, 0) and end at (1, 0). Several letters are written with a pen lift (the cross of a t or an x, the dot of an i or a j); we join the strokes of every letter end to start into a single stroke, so a few letters look unusual, but each remains a valid continuous trajectory to reproduce. In the stored trajectories the entry and exit strokes are traversed far faster than the rest of the letter (on average 10 times the median pen speed, up to 26 times). We therefore re-time every letter without changing its shape: the pen moves along the same polyline, but no segment is traversed faster than 3 times the letter’s median speed, and the result is rescaled to the same 500 ms.

Each trial follows a delayed-movement task: a cue starts one second of preparation, during which the letter’s bias ramps in and sets up a preparatory state, and a go cue withdraws the bias, after which the circuit’s own dynamics write the letter. A trial starts with 300 ms of spontaneous activity. At the cue, 1 s before the go cue, the ramp rises as 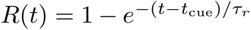 with *τ*_*r*_ = 400 ms; at the go cue it is withdrawn with a 2 ms time constant. The letter is written in the 500 ms after the go cue. During that time an arm moves the pen origin from (0, 0) to (1, 0) at constant speed, and the pen is

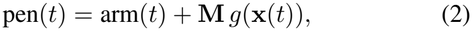

a direct linear readout of the rates, as in [10], with no filtering. The network therefore writes the letter minus the arm, which starts and ends at (0, 0).

Writing the letter is not enough: as in a real movement, the pen should stay still while the circuit prepares and come to rest when the letter ends. The learning target therefore has three parts. The letter is sampled at every integration step (every 0.5 ms, 1001 samples); the readout is also asked to hold (0, 0) during the 1 s of preparation (50 samples), so preparation is close to output-null [14], and for 500 ms after the letter (100 samples), where the pen speed is penalized as well (weight *λ* = 2 on *τ*_*s*_ **ż**, *τ*_*s*_ = 50 ms) so the pen settles smoothly at the letter’s end. We chose *λ* on a separate test (*N* = 100, letters a–d, ten draws, *λ* ∈ {0.5, 1, 2, 4}) as the largest *λ* whose mean *R*^2^ over both circuits stays within 0.002 of that at *λ* = 0.5. During preparation, and in the rest between the letters of a word, the pen is lifted, as a movement-gating (stop) signal would do; in figures of several letters a straight line joins the end of each hold to the start of the next letter.

The readout is deliberately left untrained, so that the biases do all the learning. Its weights **M**∈ ℝ^2*×N*^ are a random Gaussian pattern on the excitatory units, drawn once and never fitted to any activity or letter, times a single global gain; there is no offset. The recurrent weights are built without reference to any letter, so this global gain is the only synaptic parameter that is set with a letter in view. It is picked on the four letters a–d, with the SOC, so that their least-squares preparatory states have the 1.5 Hz spread of [10]: a choice of scale, computed in closed form, not a fit of the pen to the letters. It is shared with the MISOC of the same draw and never changed afterwards, for any letter.

### E. Learning the bias patterns

Learning a letter means finding the one bias pattern that makes the fixed circuit write it. We do this in two stages: a linear estimate that is cheap to compute, then a refinement on the full nonlinear network. Following [10], a bias pattern is parameterized by the preparatory state **a**_*l*_ it produces, **P**_*l*_ = **a**_*l*_ *−***W** *g*(**a**_*l*_), which makes **a**_*l*_ the steady state of (1) under constant drive. A first estimate solves the regularized least-squares problem

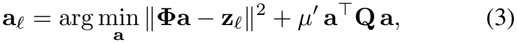

where **Φ** is the exact linear map from **a** to the pen samples (including the finite ramp and the go withdrawal), **a**^*⊤*^**Q a** is the evoked energy of [10] with **Q** scaled to unit mean eigenvalue, and *µ*^*′*^ is 0.03 times the mean eigenvalue of **Φ**^*⊤*^**Φ**; for the start, the hold-speed penalty rows are appended to **Φ** and **z**_*l*_.

The linear estimate ignores the network’s nonlinearity, so we then refine it on the real network, keeping only changes that improve the letter. Gauss–Newton steps on the full nonlinear network use the exact Jacobian of the pen samples along the simulated trajectory. A step is judged by *R*^2^ over all samples the readout is asked to match (the letter, the preparation and the hold, and the hold-speed penalty), not by the letter’s RMS error that we report; a step that does not raise it is rejected and halved, an accepted one grows by half (up to a full step), a candidate state whose spread exceeds 10 Hz is scaled down as a whole to that spread, and the search stops after 20 steps or 6 rejections in a row. The least-squares start is not capped, and it is kept when no step improves on it. Each letter is learned on its own, from rest.

We score a letter by how far the pen strays from it. Our main error measure is the root-mean-square distance between pen and target,

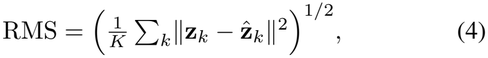

over the *K* = 1001 samples of the letter (every 0.5 ms) in the 500 ms after the go cue, given in percent of a letter width: it says directly how far the pen strays. We also report 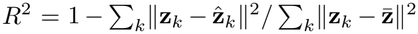. It is high for every circuit here, because it is dominated by the long strokes, and it hides differences in loops and corners that are plain to the eye.

### F. Words and return to rest

To test whether letters learned alone can be combined, we write words by playing the letters’ bias patterns one after another, without relearning, and check that the circuit settles back to rest between letters. A word is written in one continuous simulation: the bias patterns of its letters are played in order with go cues 4 s apart (1 s of preparation, the 500 ms letter, 500 ms of hold and 2 s of rest), and the arm advances one letter width per letter. Nothing is relearned. We also test each circuit’s return to rest by simulating 40 s after a single letter with no input; a circuit whose mean rate deviation is still above 0.1 Hz then has not returned to rest, and its draw (both circuits) is left out of the analyses of words and doubled letters.

### G. Draws, network size and input channels

So that no result rests on one lucky circuit, we repeat everything over many random circuits and several network sizes, and compare the SOC and MISOC built from the same draw. We use 25 independent random draws at each of 5 sizes (*N* = 50, 70, 100, 150, 200; 75/25 split): the ten seeds of our earlier sweep, plus the next seeds, in order, for which both circuits were accepted (*α <* 0.98). All seeds are frozen, so every circuit can be rebuilt exactly. At *N* = 50 one of the earlier ten MISOCs did not stabilize (*α* = 1.15) and was replaced in the same way. The readout pattern is the same for all draws of a size; each draw has its own global readout gain, set on a–d, so draws differ in their recurrent weights and in this one gain.

With one bias value per neuron, each letter has as many free numbers as the network has units. To ask how many independent signals the bias really needs, we also restrict it to **P**_*l*_ = **B c**_*l*_ with **B** ∈ ℝ^*N ×K*^ fixed, with *K* = 20 channels (and, to see how far the compression goes, *K* = 50): either *K* random input vectors or the *K* input directions that move the pen most (the top right singular vectors of the linear map from bias to pen samples, computed from the network and readout alone), scaled to unit length; the random vectors are drawn once per network size, so all draws and both circuits of a size share them. Only the *K* amplitudes **c**_*l*_ are learned.

This restricted problem needs its own learning method, because the full-bias algorithm cannot simply be reused here: run with *K* channels, by moving each state it proposes to the nearest state the channels can reach, it reproduces the full-bias results exactly when there are as many channels as units, but with fewer channels it can drive the circuit into saturated states (rate spreads of up to thousands of Hz), increasingly so in larger networks (7 of 10 circuits at *N* = 200 with 20 channels, where it also writes worse letters on average; Supplementary Section S2). We therefore learn **c**_*l*_ directly, with the same Gauss–Newton search and stopping rules but with a ridge penalty, a penalty on the sum of the squared amplitudes that keeps them from growing large, in place of the evoked energy: 0.03 times the mean eigenvalue of **L**^*⊤*^**L** for the start, where **L** maps **c**_*l*_ to all samples the readout is asked to match, and 0.03 times the mean eigenvalue of **J**^*⊤*^**J** for each step, recomputed from the current Jacobian **J**. The cap applies to the spread of the rates at the go cue, including at the start, and is enforced by rescaling **c**_*l*_, so the spread after rescaling is close to, not exactly, 10 Hz.

With few channels, a letter occasionally fails to learn, and we restart it from a weaker initial guess. The least-squares start can be too strong: it can push the circuit into a strongly nonlinear range where no step improves on it, and since only improving steps are accepted, the search then stays there. A letter whose RMS error exceeds 30% of a letter width is therefore learned again from half the least-squares start and, if it still exceeds 30%, from a quarter of it, keeping the fit with the better *R*^2^ over all samples the readout is asked to match. This was needed for 710 of the 52,000 letter fits (1.4%; 87% of them in MISOCs with random channels); 14 of them also needed the quarter start. Afterwards no letter in any input-channel test exceeds 30%. All draws enter every comparison.

## III. Results

### A. One fixed circuit, many letters

Fig. 1 shows the basic idea on the 200-unit MISOC. The letters a and b are drawn by the same network through the same readout; only the bias pattern differs. While the ramp is on, the network settles into a preparatory state with the pen lifted; when the ramp is withdrawn at the go cue, the network’s own dynamics take over and the fixed readout, added to the arm, writes the letter.

### B. The whole alphabet for free

The readout gain was set with a–d only and then frozen. We now ask the same circuits for all 52 letters, learning nothing but one bias pattern per letter (Fig. 2, Table I). No weight changes, neither recurrent nor readout, so the other 48 letters come for free as far as the circuit is concerned: a new letter is a new bias pattern. The readout is a random projection, and its one scalar was set on a–d; no weight of the circuit or the readout has ever been fitted to the other 48 letters. Both circuits write every letter recognizably (Fig. 2): over all 25 draws of both circuits at *N* = 200, no letter has an RMS error above 5.2% of a letter width. Uppercase letters, with their larger loops and crossings, are written less accurately than lowercase ones (RMS 1.0% vs 0.8% for the MISOC, 2.0% vs 1.4% for the SOC; Table I). Because each letter is learned from rest and nothing shared is adapted, adding a letter cannot change any other letter: the repertoire grows without forgetting.

**TABLE I.** Writing with *N* = 200 units (mean ± SD over 25 random circuits; words and doubled letters over the draws whose MISOC returns to rest, see text)

|  | SOC | MISOC |
|---|---|---|
| *RMS pen error (% of a letter width)* |  |  |
| All 52 letters | 1.7 ± 0.2 | 0.9 ± 0.1 |
| Uppercase / lowercase | 2.0 / 1.4 | 1.0 / 0.8 |
| Worst letter of the circuit | 3.9 ± 0.6 | 1.6 ± 0.1 |
| Doubled letter, 2nd copy | 1.7 ± 0.2 | 1.5 ± 0.6 |
| Letters in random words | 1.6 ± 0.1 | 1.4 ± 0.5 |
| 20 input channels (network-derived) | 6.3 ± 0.3 | 6.2 ± 0.4 |
| 20 input channels (random) | 7.9 ± 0.8 | 9.2 ± 1.3 |
| *R*<sup>2</sup> |  |  |
| All 52 letters | 0.996 ± 0.001 | 0.999 ± 0.000 |
| Letters with *R*<sup>2</sup> ≥ 0.95 | 100% | 100% |
| Letters in random words | 0.996 ± 0.001 | 0.997 ± 0.003 |
| 20 input channels (network-derived) | 0.946 ± 0.005 | 0.948 ± 0.009 |
| Spectral abscissa *α*(**W**) | 0.494 ± 0.005 | 0.745 ± 0.065 |
| Rate 40 s after a letter (Hz) | 0.00 ± 0.00 | 0.02 ± 0.05 |

**Fig. 2.**
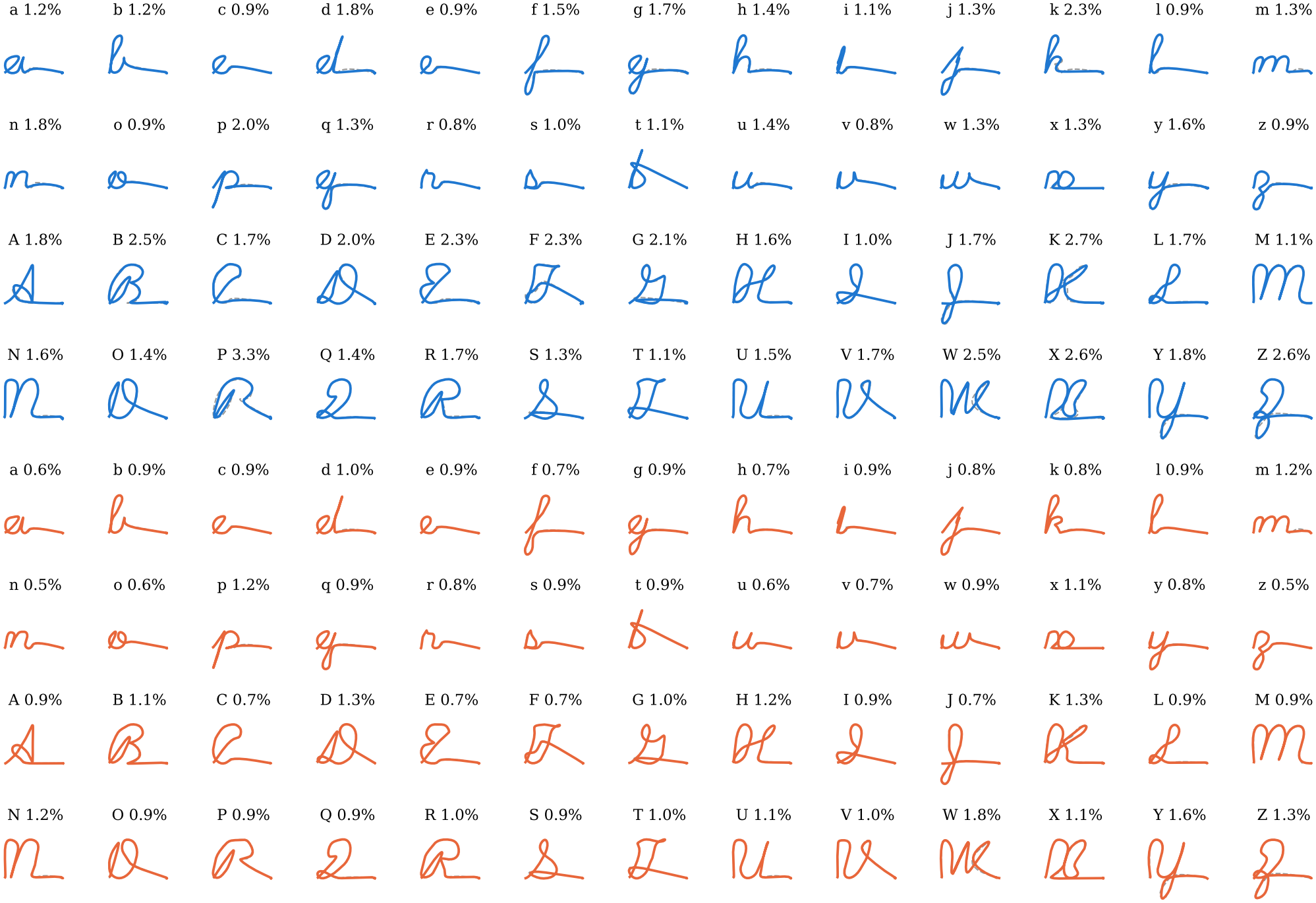
All 52 letters from bias patterns alone (*N* = 200, first draw). Top four rows: SOC; bottom four rows: MISOC. Each panel: target (dashed), pen (color) over the letter and the hold, and the RMS error in percent of a letter width. The readout was set with a–d and never changed.

### C. Mutual inhibition makes the circuit write better

Replacing the SOC’s individually tuned inhibitory-to-inhibitory synapses with one shared weight makes the circuit write better, although fewer of its weights are optimized. Over 25 random circuits of 200 units, the MISOC’s pen stays on average 0.9% of a letter width from the target and the SOC’s 1.7%: the MISOC makes 48% less error, and does so in 25/25 circuits (paired Wilcoxon *p <* 0.001; Table I). The difference is visible in the letters themselves (Fig. 2), and is largest where it matters most: the worst letter of a circuit strays on average 1.6% for the MISOC and 3.9% for the SOC. By *R*^2^ both circuits look nearly perfect (0.999 and 0.996; 100% and 100% of letters at *R*^2^*≥* 0.95; MISOC better in 25/25 circuits, *p <* 0.001). The advantage holds at every network size: from *N* = 200 down to *N* = 50, the MISOC writes with significantly less error than the SOC at 5 of 5 sizes, 16– 48% less, and the advantage grows with network size (Fig. 3b, Table II). It also survives compressing the bias to 50 network-derived input channels (Section III-F). The next section shows that this benefit needs the mutual inhibition to be moderate.

**TABLE II.** Network size: RMS pen error (percent of a letter width) and *R*^2^, means over 52 letters and 25 circuits per size, with paired wilcoxon *p* for the RMS error of MISOC vs SOC.

|  | one bias per unit |  |  |  |  | 20 input channels |  |  |
|---|---|---|---|---|---|---|---|---|
|  | RMS (%) |  |  | *R*<sup>2</sup> |  | RMS (%) |  |  |
| *N* | SOC | MISOC | *p* | SOC | MISOC | SOC | MISOC | *p* |
| 200 | 1.7 | 0.9 | <0.001 | 0.996 | 0.999 | 6.3 | 6.2 | 0.508 |
| 150 | 1.9 | 1.1 | <0.001 | 0.995 | 0.998 | 6.2 | 6.2 | 0.874 |
| 100 | 2.4 | 1.7 | <0.001 | 0.992 | 0.996 | 6.5 | 6.7 | 0.052 |
| 70 | 2.9 | 2.3 | <0.001 | 0.988 | 0.993 | 6.3 | 6.7 | 0.071 |
| 50 | 3.4 | 2.8 | <0.001 | 0.984 | 0.989 | 6.6 | 7.7 | <0.001 |

**Fig. 3.**
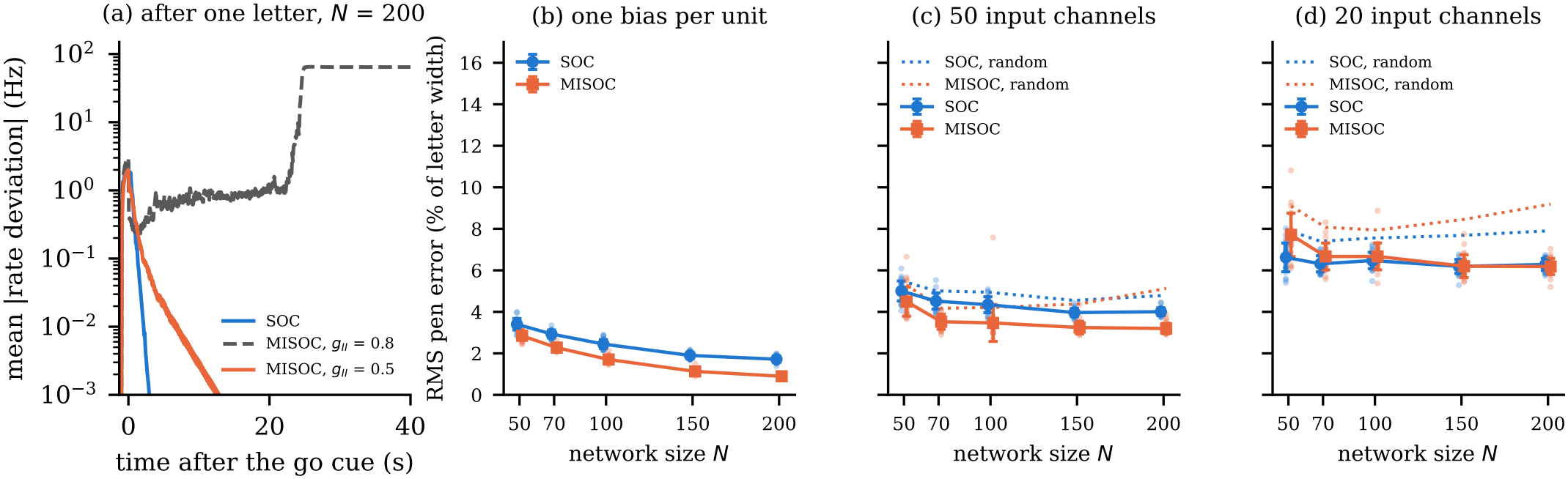
(a) Return to rest: mean rate deviation after one letter (a), *N* = 200, for the SOC, the earlier MISOC with *g*_*II*_ = 0.8 and the MISOC with *g*_*II*_ = 0.5. (b–d) Network size, 25 random circuits per size, RMS pen error in percent of a letter width (mean *±*SD; dots: circuits): (b) one bias value per unit; (c) 50 and (d) 20 fixed input channels, the network’s top input directions (solid) or random channels (dotted). Each circuit has its own readout gain, set on a–d.

### D. Mutual inhibition must be moderate

Mutual inhibition helps only when it is moderate (Fig. 3a). In our earlier circuits the shared weight was *g*_*II*_ = 0.8 (these numbers and the *g*_*II*_ = 0.8 trace in Fig. 3a come from our earlier simulations, which used a 50 ms readout filter, the stored letter timing and ten draws); those MISOCs wrote single letters well, but at *N* = 200 every one of them (10/10 draws) ran away to a saturated state (mean rate deviation *≈*74 Hz) within 5–40 s of a single letter, and a letter written right after another was badly distorted (*R*^2^ = 0.46 for the second copy of a doubled letter). With *g*_*II*_ = 0.5 the MISOC returns to rest like the SOC (0.02 Hz 40 s after a letter; SOC 0.00 Hz) and the second copy of a doubled letter is written well (RMS 1.5% vs 0.9%; *R*^2^ 0.996 vs 0.999). Across all sizes, 110 of the 125 MISOCs with *g*_*II*_ = 0.5 settle completely (below 0.1 Hz 40 s after a letter), 6 keep a small residual (0.2–0.4 Hz) but still write words, and 9 (*N* = 50, 70, 100) keep residual activity (0.6–5.1 Hz) that disrupts the letters that follow; 125/125 SOCs settle. We leave the draws of the 15 MISOCs that do not settle (12%) out of the analyses of doubled letters and words.

The MISOC’s spectral abscissa is higher than the SOC’s (at *N* = 200, 0.745 against 0.494), a consequence of the shared weight, which holds the inhibitory difference modes near +*g*_*II*_. Its advantage is not due to this. Stopping each SOC’s own optimization as soon as its abscissa reaches that of the MISOC of the same draw leaves its writing essentially unchanged (*N* = 100, first ten draws: RMS 2.4% against 2.4% for the fully optimized SOC, *p* = 0.32), and the MISOC (1.7%) still writes better than the matched SOC in 10 of the 10 draws (Supplementary Section S1).

### E. Words from letters learned alone

Because the circuits return to rest, the pen settles during the hold and is lifted during the rest and the preparation, letters learned one at a time can be strung together. We wrote ten random five-letter words with mixed case, each in one continuous run (Fig. 4). Over the draws whose MISOC returns to rest, the letters inside words stray on average 1.4% of a letter width for the MISOC and 1.6% for the SOC (MISOC better in 17/23 circuits, *p* = 0.020; *R*^2^ 0.997 and 0.996); no word was ever used for learning.

**Fig. 4.**
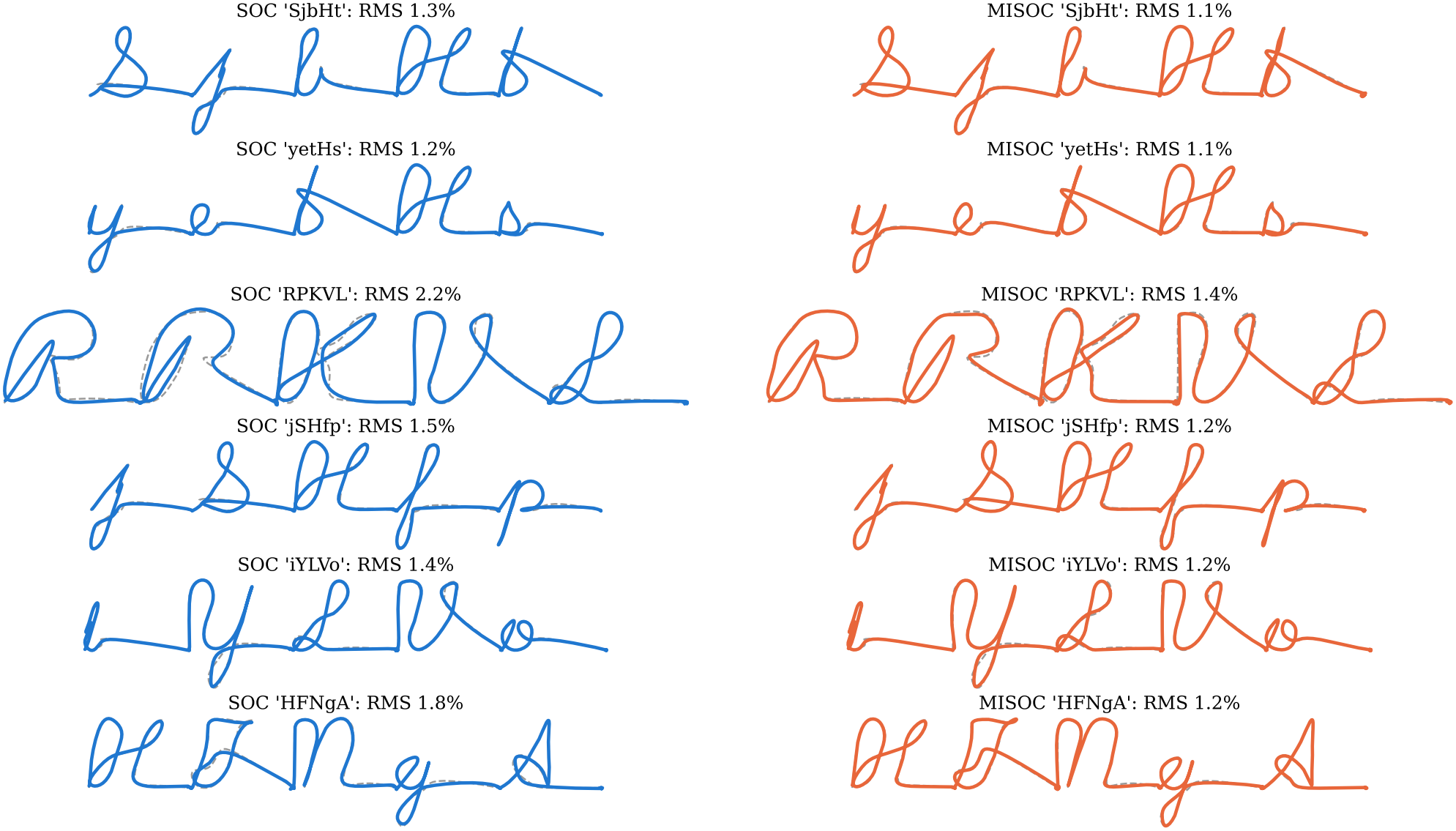
Six of the ten random words, each written in one continuous run from letters learned one at a time (*N* = 200, first draw; left: SOC, right: MISOC; dashed: targets). The pen path of each letter and its hold is drawn, and a straight line joins each hold to the next letter, across the rest and the preparation, in which the pen is lifted. The value is the mean RMS error of the word’s letters, in percent of a letter width.

### F. How small, and how few signals?

Circuits as small as 50 units write the alphabet, and the MISOC keeps its advantage at every size (Section III-C). A letter also does not need one bias value per neuron. With 20 fixed input channels and only 20 learned amplitudes per letter (Fig. 3d), the 200-unit MISOC writes the alphabet with an RMS error of 6.2% using the network’s own top input directions (*R*^2^ = 0.948) and 9.2% using random channels (SOC: 6.3% and 7.9%); the MISOC is better in 13/25 circuits (*p* = 0.508). With so few channels the advantage of the MISOC disappears, but it does not reverse: the two circuits do not differ significantly at *N* = 200, 150, 100 and 70 (the SOC is slightly ahead at 100 and 70), and only at *N* = 50, where twenty channels must drive a circuit of just 50 units, is the SOC significantly more accurate (Table II). This marks a limit on how far a small MISOC can be compressed, not a flaw of mutual inhibition. With random channels the SOC is more accurate at every size (Table III). The letters remain recognizable but lose accuracy, because each letter must now also park the pen during preparation and the hold with only 20 amplitudes.

**TABLE III.**
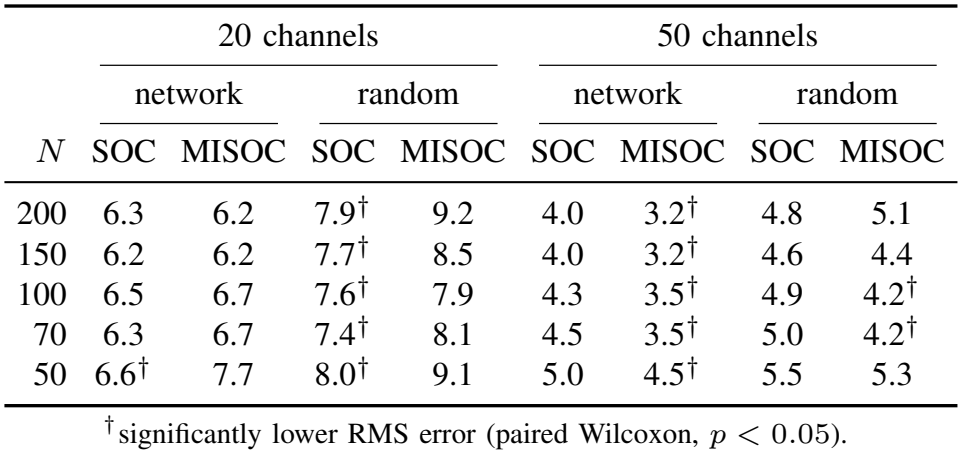
Input channels: RMS pen error (percent of a letter width), mean over 52 letters and 25 circuits per size, with 20 or 50 fixed input channels. *Network:* the network’s top input directions; *random*: random vectors.

|  | 20 channels |  |  |  | 50 channels |  |  |  |
|---|---|---|---|---|---|---|---|---|
|  | network |  | random |  | network |  | random |  |
| *N* | SOC | MISOC | SOC | MISOC | SOC | MISOC | SOC | MISOC |
| 200 | 6.3 | 6.2 | 7.9<sup>†</sup> | 9.2 | 4.0 | 3.2<sup>†</sup> | 4.8 | 5.1 |
| 150 | 6.2 | 6.2 | 7.7<sup>†</sup> | 8.5 | 4.0 | 3.2<sup>†</sup> | 4.6 | 4.4 |
| 100 | 6.5 | 6.7 | 7.6<sup>†</sup> | 7.9 | 4.3 | 3.5<sup>†</sup> | 4.9 | 4.2<sup>†</sup> |
| 70 | 6.3 | 6.7 | 7.4<sup>†</sup> | 8.1 | 4.5 | 3.5<sup>†</sup> | 5.0 | 4.2<sup>†</sup> |
| 50 | 6.6<sup>†</sup> | 7.7 | 8.0<sup>†</sup> | 9.1 | 5.0 | 4.5<sup>†</sup> | 5.5 | 5.3 |

<sup>†</sup>significantly lower RMS error (paired Wilcoxon, *p* < 0.05).

Fifty channels show where the limit lies (Fig. 3c, Table III). With 50 network-derived channels the MISOC again writes with significantly less error than the SOC at 5 of 5 sizes (10– 22% less; at *N* = 200, 3.2% against 4.0%). With 50 random channels the MISOC is ahead at 2 of 5 sizes (15–17% less) and behind at none. At *N* = 50, fifty channels are as many as there are units.

### G. The whole alphabet in one stroke

As a last test, Fig. 5 writes a–z and A–Z, each as one continuous run of 26 letters with the 200-unit MISOC (RMS error 1.1% and 1.3% of a letter width; worst letter 1.5% and 2.1%). The SOC does the same with 1.3% and 1.9% (worst letter 2.3% and 3.3%). Each letter was learned alone; the arm and the settling pen join them.

**Fig. 5.**
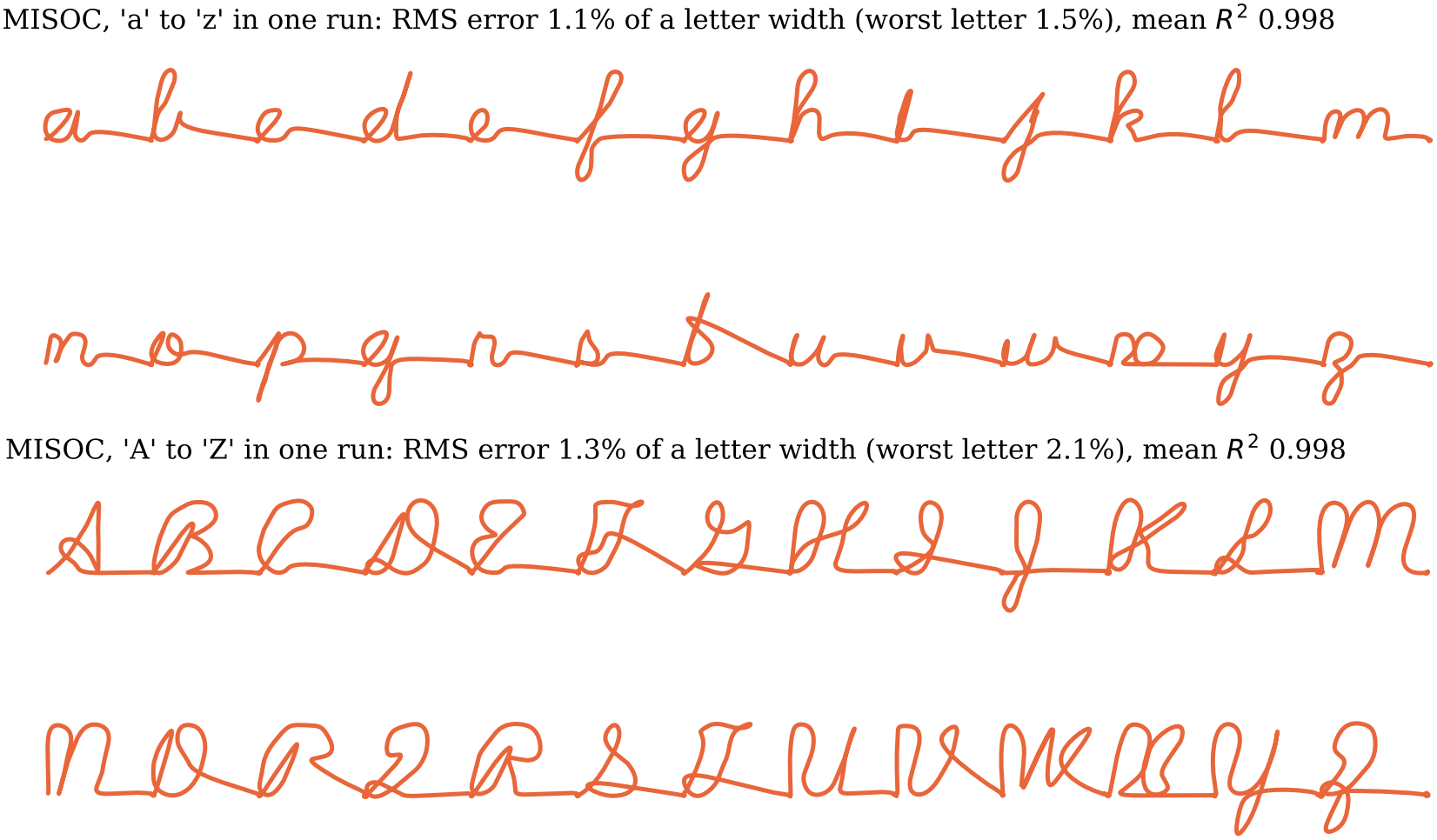
a–z and A–Z, each written in one continuous run of 26 letters by the MISOC (*N* = 200, first draw). Each run is continuous; it is drawn on two lines, broken between m and n, only to fit the page. A straight line joins each letter’s hold to the next letter, across the rest and the preparation, in which the pen is lifted. Letters were learned one at a time and never together.

## IV. Discussion

### A. What is new

In these circuits the identity of a letter lives entirely in a bias pattern. The recurrent weights and the readout are shared by all 52 letters, and the readout is a random projection whose only free quantity was set on four letters. A fixed circuit thus acts as a library of movements addressed by preparatory bias. None of the ingredients is new on its own, and we did not set out to reinvent them. The dynamics are those of Hennequin *et al*. [10]; preparatory states as the selector of movements come from motor cortex [1], [2]; learning only biases is known to be expressive [8]; and in our own work, bias currents alone switch a four-neuron circuit between 24 logic functions [6], while mutual inhibition between inhibitory neurons enlarges the set of functions a small circuit can reach [7]. What this study adds is the connection: reading the preparatory input as a bias joins these threads, and a fixed circuit then becomes a store for a growing repertoire, in which each movement costs one input pattern, adding one never disturbs the others, and 50 to 200 units hold the whole alphabet. Four results follow from that connection.

First, the repertoire. One fixed, sparse circuit of plain rate units writes 52 two-dimensional cursive letters, each selected by nothing but its bias pattern, and 48 of them were never seen by any weight. Earlier fixed-weight models driven by preparatory or tonic input produced a handful of movements: two drawn trajectories [10], one-dimensional traces [11] or reaches [12], [13]; the closest model that wrote new letters from a learned input first trained an LSTM, with about 80 000 weights by our estimate, on letters [9].

Second, the whole movement is accounted for, not only its start. The bias is an input the circuit receives, the preparatory state is reached during a second of preparation while the pen is held close to still, and in most circuits the activity comes back to rest after the letter. This is what lets letters, each learned alone and from rest, chain into words and whole alphabets in one continuous run with nothing relearned.

Third, continual learning comes for free. A new letter is a new bias pattern and nothing else: no recurrent weight and no readout weight is touched, so learning it cannot disturb any letter already learned, and the repertoire can grow letter by letter without replay, weight consolidation or retraining on old letters. Interference between old and new skills, catas-trophic forgetting, is the central problem of continual learning in trained networks; here it cannot arise, by construction. Together with the chaining above, a repertoire learned one movement at a time is also usable in any order.

Fourth, the circuit’s structure decides how well bias can control it. Replacing the SOC’s individually tuned inhibitory-to-inhibitory synapses with one shared weight removes parameters and makes the circuit more accurate at every size, and the advantage survives compressing the bias to fifty network-derived input channels. Bias control is therefore not a generic property of any stable recurrent network that one can take for granted; it depends on how inhibition is organized, as it does for logic in small circuits [7]. Carried from discrete logic to continuous, high-dimensional motor output, this makes bias-controlled reconfigurability a candidate mechanism for holding a large and growing motor repertoire in a fixed circuit.

### B. Relation to previous models

Our circuits build on those of Hennequin *et al*. [10], but we invert what is learned. They fixed the preparatory states, choosing the two most amplified initial conditions of the SOC, set each input pattern by formula so that the network reaches its state, and learned the readout weights, with an offset, by least-squares regression so that the activity traced two target movements. They never learned a bias. Here a bias is learned for every letter and the readout is not: its weights are drawn at random and never fitted, and only a single global gain, one scalar number, is chosen, on a–d.

Williams *et al*. [8] proved that random networks with learned biases are universal approximators and trained recurrent networks to produce simple trajectories, one per network. Here one network holds 52 movements, and its bias acts before the movement and is withdrawn at its start.

Stroud *et al*. [11] found that gain changes, but not a trained tonic input that stayed on during the movement, produced new movements from a fixed SOC. As a baseline for their gain method they also trained a separate initial condition for each output and found it about as effective. That test was a single panel: ten 1-D traces, each learned with the initial state placed directly into the network at movement onset. In it, as in their gain experiments, the readout was fitted first: all of its weights, and an offset, were learned by least squares so that the untrained network produced an initial movement, and were then held fixed while the gains or initial conditions were learned. Our readout weights are never fitted: they are drawn at random, and only their single global gain, one scalar number, is chosen. The test also does not ask whether the state can be reached by an input the circuit receives, what the output does while the circuit gets there and after the movement ends, whether separately learned movements can follow one another, or how the circuit’s structure matters. Here the bias is a preparatory ramp withdrawn at the go cue, a way of setting that initial condition: it chooses where the circuit starts, and the circuit’s own dynamics do the rest, as preparatory activity is thought to do in motor cortex [1], [2], [12]. Optimal preparatory inputs to fixed networks have been studied for a few reaches [12], [13]. In all of this work the preparatory input drives simple outputs, a few reaches or one-dimensional traces, nothing approaching 52 two-dimensional cursive letters; and here the readout’s single global gain is picked on a–d alone, so 48 of the letters are written by a readout never fitted to them.

The closest machine-learning result uses far more machinery. Fabi *et al*. [9] trained an LSTM, whose units each carry an internal memory and multiplicative gates, on thirteen letters, with 440 handwritten samples per letter, and then wrote further letters by adapting only the weights into its input layer, which, with a constant one-hot label, amounts to learning a constant input of 100 values to its frozen recurrent core. That input stays on throughout the movement and, through the gates, can reshape the network’s dynamics as it unfolds, and the LSTM alone has about 80 000 trained weights (our estimate from the reported architecture, one LSTM layer of 100 units with 100 inputs). Our network is a generic, sparse recurrent network of plain rate units that is never trained on letters: our largest circuits have about 6 800 (SOC) and 8 300 (MISOC) fixed synapses, and at *N* = 100 the SOC has about 1 700 and the MISOC about 2 100, some 17–21% of all possible connections, with the MISOC’s 600 inhibitory-to-inhibitory synapses sharing a single weight, *g*_*II*_. Every letter, including the first, is learned from a single sample trajectory, the input is withdrawn before the movement begins, and it can be compressed to as few as 20 values per letter. Duan *et al*. [17] copied drawings made of simple segments with modular recurrent networks trained end to end.

Logiaco *et al*. [18] chained motor motifs without interference, using a 500-unit cortical network and a thalamocortical loop of synapses learned for every motif. Their motifs are simple: each is a single scalar output, read out linearly from the network, such as a sinc function or a slow oscillation. Here a sparse circuit of 50 to 200 plain rate units, with nothing learned but one bias pattern per letter, writes words and whole alphabets of two-dimensional cursive letters in one continuous run; to our knowledge no other model with fixed weights has strung together movements of this complexity.

### C. Continual learning by construction

Learning a new letter changes only that letter’s bias pattern, so it cannot interfere with any letter already learned; nothing like replay or weight consolidation is needed. The same holds for movement-specific gain patterns [11], and for the per-motif loops of Logiaco *et al*. [18]; here the only per-motif quantity is an input. With moderate mutual inhibition and a pen that settles, letters learned alone also compose into words in one continuous run. The price is storage: each letter is an input pattern. With 20 channels that pattern has 20 numbers, about twice the dimension of the letters themselves (the 52 letters span ten dimensions at 95% of their variance), and the alphabet is still written recognizably (RMS error *≈*6.2%), though less accurately than with one bias per neuron.

### D. Mutual inhibition

Connections between inhibitory neurons are abundant in cortex [19], [20], and even more common in human than in mouse cortex [21]. Replacing the finely tuned inhibitory-to-inhibitory block of the SOC with one shared all-to-all weight removes parameters and, at a moderate strength, improves writing. The strength matters: at *g*_*II*_ = 0.8 the inhibitory difference modes sit close to instability, and the circuit, linearly stable at rest, is pushed into runaway activity by its own movements. Mutual inhibition expands what small circuits can do [7]; it also needs to stay moderate. The benefit needs a rich enough drive: with one bias per neuron the MISOC writes with 16–48% less error than the SOC, and with fifty network-derived input channels it still has the advantage at every size, whereas with twenty channels the advantage is largely gone. Twenty channels are thus near the limit of how far the bias of these circuits can be compressed. We read this as a boundary on the mechanism rather than a failure of it. The SOC’s per-synapse inhibitory block lets any single inhibitory unit be nudged without moving the rest; the MISOC’s shared weight ties them together, so that they move as a group, and the circuit needs enough independent preparatory inputs to exploit that coupling, and twenty channels are too few.

### E. Limitations

The circuits are noise-free rate networks. *R*^2^ is lenient for small features: a letter at *R*^2^ *≈* 0.94 can still show a visibly misshapen loop, which is why we lead with the RMS error. The letters were re-timed to remove the near-instantaneous entry and exit strokes of the stored trajectories; the shapes are unchanged, but the speed profile is ours. The MISOC needs more stability-optimization steps than the SOC to be built, small MISOCs are often hard to stabilize at all (at *N* = 50, 45 of the 70 MISOCs we built were not accepted), and we have not mapped how accuracy and stability depend on *g*_*II*_ beyond 0.5 and 0.8. Some MISOCs (12%) do not return to rest after a letter, and with few input channels a few letters are learned only from a weaker start (Methods). Finding a construction or learning rule that reliably yields MISOCs that settle is future work; since inhibitory interneurons are densely and mutually connected in cortex [19], [20], one would expect biology to have a solution. Bias patterns were found by gradient methods with exact Jacobians, not by a biologically plausible rule.

## V. Conclusion

A stability-optimized circuit with fixed recurrent weights and a fixed random readout writes all 52 cursive letters when, and only when, it receives the right bias pattern. A readout calibrated on four letters serves the whole alphabet, new letters never disturb old ones, letters learned alone chain into words, and moderate mutual inhibition makes circuits significantly more accurate at every size we tested, from 50 to 200 units, with most circuits coming back to rest. Bias currents deserve a place alongside synaptic plasticity as a way for neural circuits, and for the hardware that imitates them, to hold many behaviors at once.

## Acknowledgment

The authors thank Ching-Che Charng for valuable editing advice and discussions of this paper.

An AI assistant (Claude, Anthropic) was used to translate the authors’ pseudocode into simulation code, to write the figure-generation code and to help draft and edit the manuscript. All ideas, the study design and the interpretation of the results are the authors’. The authors checked every line of the code and the text and take full responsibility for the content.

## Notes

This work was supported by the U.S. Army Research Office, DEVCOM Army Research Laboratory, through a Short-Term Innovative Research (STIR) grant, Project W911NF261A006, and by the National Science and Technology Council, Taiwan, under grant NSTC 114-2321-B-007-006. The views and conclusions contained in this paper are those of the authors and should not be interpreted as representing the official policies, either expressed or implied, of the Army Research Office or the U.S. Government. The U.S. Government is authorized to reproduce and distribute reprints for Government purposes notwithstanding any copyright notation herein.

### Competing Interest Statement

The authors have declared no competing interest.

