## Supplement for "Preparatory Bias Currents Alone Write Letters and Words in a Fixed Recurrent Circuit"

### S1. The MISOC's higher abscissa does not explain its advantage

The MISOC cannot reach as low a spectral abscissa as the SOC: its shared inhibitory-to-inhibitory weight holds the inhibitory difference modes near $+g_{II}$ (main text, Sec. II-C), so its optimization stops near $\alpha = 0.6$ or above, while the SOC's stops below 0.5. A circuit closer to instability amplifies more, so the MISOC's better writing could in principle come from its abscissa rather than from its mutual inhibition.

To separate the two, we rebuilt the SOC of each of the first ten draws at $N = 100$ from the same initial matrix, with the same optimization, but stopped it as soon as its abscissa fell below that of the MISOC of the same draw (`v2/build_socam.py`). All 52 letters were then learned on these abscissa-matched SOCs exactly as for the other circuits, with the draw's readout. The matched SOCs write like the fully optimized ones (mean RMS error 2.40% against 2.42%, $p = 0.32$, paired Wilcoxon over the ten draws), and the MISOC (1.67%) writes better than the matched SOC in all ten draws ($p = 0.002$; Table S1). Stopping the optimization early thus raises the SOC's abscissa to the MISOC's without improving its writing.

TABLE S1
Abscissa-matched SOCs ($N = 100$, first ten draws). Spectral abscissa of the fully optimized SOC, of the SOC stopped at the MISOC's abscissa (matched), and of the MISOC, and the RMS pen error of each over all 52 letters (percent of a letter width).

| | spectral abscissa $\alpha$ | | | RMS error (%) | | |
|---|---|---|---|---|---|---|
| seed | SOC | matched | MISOC | SOC | matched | MISOC |
| 1 | 0.485 | 0.589 | 0.600 | 2.41 | 2.40 | 1.47 |
| 7 | 0.489 | 0.587 | 0.600 | 2.22 | 2.14 | 1.64 |
| 9 | 0.500 | 0.576 | 0.600 | 2.12 | 2.13 | 1.62 |
| 12 | 0.468 | 0.672 | 0.681 | 2.37 | 2.36 | 1.70 |
| 16 | 0.477 | 0.713 | 0.714 | 2.52 | 2.51 | 1.71 |
| 17 | 0.480 | 0.555 | 0.600 | 2.67 | 2.67 | 1.88 |
| 18 | 0.480 | 0.590 | 0.600 | 2.37 | 2.36 | 1.56 |
| 20 | 0.478 | 0.598 | 0.600 | 2.20 | 2.22 | 1.68 |
| 28 | 0.476 | 0.595 | 0.600 | 2.39 | 2.43 | 1.71 |
| 29 | 0.483 | 0.589 | 0.600 | 2.90 | 2.83 | 1.76 |
| mean | | | | 2.42 | 2.40 | 1.67 |

### S2. Why the input-channel test has its own learning method

With one bias value per unit, the bias pattern of a letter is learned through the preparatory state $\mathbf{a}$ it produces (main text, Sec. II-E): Gauss–Newton steps on $\mathbf{a}$, regularized by the evoked energy, with the spread of $\mathbf{a}$ capped at 10 Hz. When the bias is restricted to $K$ fixed input channels, $\mathbf{P} = \mathbf{B}\,\mathbf{c}$, only the states that the channels can hold are reachable. The most direct way to keep the full-bias method is to run it unchanged on the state and move every state it proposes to the nearest reachable one: the amplitudes $\mathbf{c}$ whose steady state $\mathbf{a}(\mathbf{c})$, solving $\mathbf{a} - \mathbf{W}g(\mathbf{a}) = \mathbf{B}\mathbf{c}$, is closest to the proposal (Gauss–Newton on $\mathbf{c}$).

*When there are as many channels as units, this is the full-bias method.* At $N = K = 50$ every state is reachable, and over all 50 circuits (25 draws, SOC and MISOC) every one of the 2600 letters matches the full-bias result exactly (Table S2, first row).

*With fewer channels it breaks down.* A proposed state is generally not reachable, and the reachable state nearest to it can lie on a saturated branch of the circuit: the strongly non-normal SOCs amplify the part of the drive that the channels cannot deliver. Some letters then end in states with rate spreads of thousands of Hz (Table S2, Fig. S1). The breakdown grows with network size: it affects 3 of 50 circuits at $N = 50$ with 20 channels, but 4 of 10 at $N = 100$ with 50 channels and 7 of 10 at $N = 200$ with 20 channels. At $N = 200$ it also raises the mean error, from 6.2% to 7.5% of a letter width; at the two smaller sizes its mean error is lower than that of the method of the main text, because most of its letters are unaffected. Because one method must serve every network size, and this one drives most large circuits into saturation, the input-channel test learns the amplitudes $\mathbf{c}$ directly (main text, Sec. II-G), with the same Gauss–Newton search and stopping rules, a ridge penalty in place of the evoked energy, the cap on the spread of the rates at the go cue, and a restart from a weaker start for letters that fail (Sec. S3-J).

TABLE S2
The full-bias algorithm run with $K$ network-derived input channels. Reference: the full-bias method for $K = N$; the method of the main text otherwise. The full-bias method caps the spread at 10 Hz, so a larger spread marks a state the channel projection has driven past the cap.

| $N$ | $K$ | circuits | letters with spread $>$ 10 Hz | circuits with such a letter | largest spread (Hz) | mean RMS (%) full-bias alg. | reference |
|---|---|---|---|---|---|---|---|
| 50 | 50 | 50 | 0.0% | 0/50 | 10.0 | 3.13 | 3.13 |
| 50 | 20 | 50 | 0.1% | 3/50 | 4,573.6 | 6.67 | 7.17 |
| 100 | 50 | 10 | 1.7% | 4/10 | 1,800.0 | 4.07 | 4.42 |
| 200 | 20 | 10 | 5.6% | 7/10 | 117,484.3 | 7.45 | 6.18 |

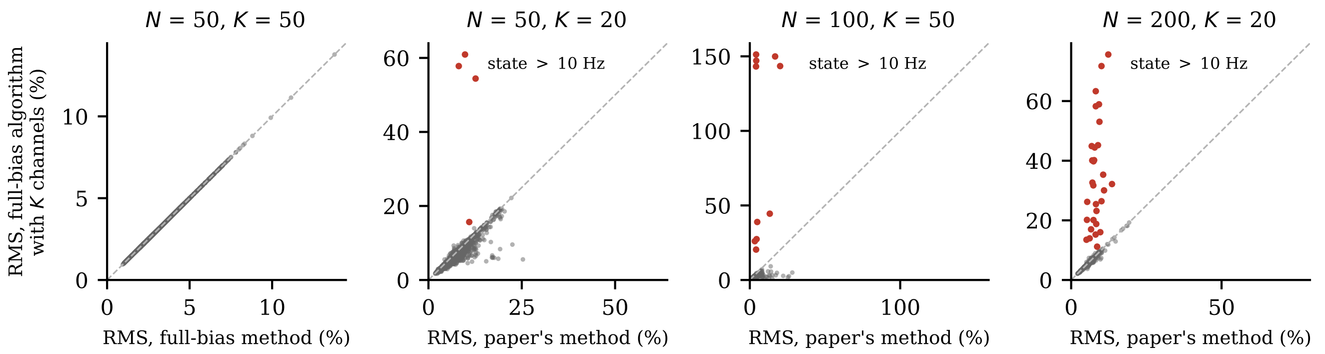


Fig. S1. Letter by letter: RMS error of the full-bias algorithm run with $K$ channels against the reference method (dashed: equal). Red: letters whose state ends with a rate spread above 10 Hz.

### S3. Supplementary Methods

This section describes the pipeline as the code implements it, in enough detail to rebuild every number in the paper. It also lists choices the main text states only briefly, and places where the implementation differs from a literal reading of the main text. Code names are in `typewriter`; paths are relative to the repository root.

#### A. Code map and reproduction

Table S3 maps the scripts to their outputs. The model code is in `soc_common/` (`rate_model.py`, `soc.py`, `learn.py`, `task.py`, `letters.py`, `config.py`). The paper pipeline is in `v2/`; the paper's results come from `v2/final.py` run with the environment settings `FINAL_ACAP=10` (the 10 Hz spread cap), `FINAL_TAG=_rt25_cap10` (output folder) and `FINAL_GAP=2` (2 s of rest between the letters of a word). Every job is cached per draw, circuit and letter set, so an interrupted run resumes where it stopped. The simulations are deterministic: rerunning a job reproduces the stored result to within $10^{-11}$. `v2/quick_check.py`, run with the same settings, checks this by relearning letters on three frozen circuits and comparing them with the stored results.

#### B. Network and integration

The potentials obey Eq. (1) of the main text, integrated with forward Euler steps of $\Delta t = 0.5$ ms:

$$\mathbf{x}_{n+1} = \mathbf{x}_n + \tfrac{\Delta t}{\tau}\big( - \mathbf{x}_n + \mathbf{W} g(\mathbf{x}_n) + R(t_n)\, \mathbf{P}_\ell \big), \qquad (1)$$

with $\tau = 200$ ms. Rest is $\mathbf{x} = 0$, where every unit fires at the 5 Hz baseline. It is a fixed point because $g(0) = 0$, and every trial starts there, 300 ms before the cue. Since $g'(0) = 1$, the linearization at rest is $\tau\dot{\mathbf{x}} = (\mathbf{W} - \mathbf{I})\mathbf{x}$, so rest is stable when $\alpha(\mathbf{W}) < 1$. Rates are recorded at every Euler step. The pen is the unfiltered readout $\mathbf{M}g(\mathbf{x})$ plus the arm. The rate model's Ornstein–Uhlenbeck noise is switched off for learning and for every result in the paper (Sec. S3-K).

TABLE S3
Scripts and what they produce.

| Script | Output |
|---|---|
| `soc_common/experiments.py` | first 10 draws per size (seed search with SOC and the earlier $g_{II} = 0.8$ MISOC) |
| `v2/build05.py` | MISOC with $g_{II} = 0.5$ for each draw |
| `v2/replace_draw.py` | replacement of a draw whose MISOC is not accepted |
| `v2/build25.py` | draws 11–25 per size |
| `v2/run_retime_ad.sh`, `select_lambda.py` | choice of $\lambda$ |
| `v2/final.py learn` | bias patterns, per-letter scores, return to rest |
| `v2/final.py evaluate` | doubled letters, random words |
| `v2/final.py lowdim` | input-channel test ($K = 20, 50$) |
| `v2/refix_lowdim.py` | the restart of failed letters (Sec. S3-J), applied to results computed before `final.py` included it |
| `v2/build_socam.py`, `final.py learn --circuits socam`, `am_report.py` | Section S1 |
| `v2/final.py lowdim_matched`, `check_reach.py`, `supp_reach.py` | Section S2 |
| `v2/final.py noise` | robustness to noise (not in the paper) |
| `v2/paper_final.py` | all numbers, tables and data figures of the paper |
| `v2/quick_check.py` | reproduction check |

#### C. Circuit construction

*Initial matrix.* Let $N_E = \mathrm{round}(0.75N)$ and $N_I = N - N_E$. Every entry exists with probability $p = 0.1$. Excitatory weights are $w_0/\sqrt{N}$ and inhibitory weights $-\gamma w_0/\sqrt{N}$, with $\gamma = 3$ and $w_0 = R/\sqrt{p(1-p)(f_E + f_I\gamma^2)}$, where $R = 10$ and $f_E$, $f_I$ are the population fractions. $W_{ij}$ is the weight from unit $j$ to unit $i$.

*Free synapses.* The E→E block never changes. The existing E→I synapses are tuned but never created. Each inhibitory synapse (I→E, and I→I in the SOC) that does not already exist is given a free slot with probability $(0.4 - p)/(1 - p)$, starting at zero weight. Up to 40% of each inhibitory block can therefore carry weight.

*Optimization.* Each step works on $\mathbf{A} = \mathbf{W} - s\mathbf{I}$, with $s = \max(1.5\,\alpha, \alpha + 0.2)$. It solves the two Lyapunov equations $\mathbf{A}^\top\mathbf{Q} + \mathbf{Q}\mathbf{A} = -2\mathbf{I}$ and $\mathbf{A}\mathbf{P} + \mathbf{P}\mathbf{A}^\top = -2\mathbf{I}$. The gradient of the smoothed spectral abscissa is $\mathbf{QP}/\mathrm{tr}(\mathbf{QP})$, and the free weights move by $-10$ times it. Then three corrections follow:

1) Dale's law: weights of the wrong sign are clipped to zero.
2) Each tuned block is rescaled to its fixed mean. The E→I mean stays at its initial value; the I→E mean is $-\gamma$ times the E→E mean; in the SOC, the I→I mean is $-\gamma$ times the E→I mean.
3) A free inhibitory synapse whose weight is exactly zero gives its slot to a random unused slot in the same row.

In the MISOC the I→I block is $-g_{II}$ off the diagonal and zero on it, and never changes.

*Stopping and acceptance.* The SOC optimizes until $\alpha < 0.5$. If it has not reached that after 4000 steps, it runs for up to 8000 more, stopping as soon as $\alpha < 0.98$. The MISOC ($g_{II} = 0.5$) optimizes until $\alpha < 0.6$, for at most 20 000 steps. A circuit is accepted when $\alpha < 0.98$.

Table S4 gives the resulting abscissae. Every SOC stopped at its target ($\alpha \leq 0.50$). Most MISOCs never reached 0.6: they ran the full 20 000 steps and ended with $\alpha$ up to 0.97. The two circuits of a draw therefore differ in their abscissa as well as in their I→I block; at $N = 200$, mean $\alpha$ is 0.49 for the SOC and 0.75 for the MISOC. Both start from the same initial matrix (same seed). Their slot assignment and rewiring, however, follow different random sequences, because the SOC also draws I→I slots.

TABLE S4
SPECTRAL ABSCISSA OF THE 25 CIRCUITS PER SIZE, AND HOW MANY MISOCS RAN THE FULL 20 000 STEPS WITHOUT REACHING $\alpha < 0.6$.

| $N$ | SOC mean (range) | MISOC mean (range) | MISOC at limit |
|---|---|---|---|
| 50 | 0.48 (0.45–0.50) | 0.74 (0.60–0.97) | 15 |
| 70 | 0.48 (0.45–0.50) | 0.66 (0.60–0.93) | 11 |
| 100 | 0.48 (0.44–0.50) | 0.71 (0.60–0.97) | 14 |
| 150 | 0.49 (0.47–0.50) | 0.68 (0.60–0.90) | 20 |
| 200 | 0.49 (0.49–0.50) | 0.75 (0.61–0.89) | 25 |

*Which draws are used.* The draws are frozen seeds that passed acceptance, taken in seed order. The first ten per size come from our earlier sweep, which accepted a seed when the SOC and the earlier MISOC ($g_{II} = 0.8$) were both stable. The other fifteen are the next seeds for which the SOC and the $g_{II} = 0.5$ MISOC were both accepted. One draw at $N = 50$ was replaced by seed 10, because its rebuilt MISOC did not stabilize ($\alpha = 1.15$).

Seeds were rejected almost only because the MISOC failed:

- Earlier sweep: 21 of 31 seeds at $N = 50$, 7 of 17 at 70, 20 of 30 at 100, 3 of 13 at 150 and 11 of 21 at 200.
- Extension to 25 draws: 54 of 69 seeds at $N = 50$ (9 of them because the SOC failed), 4 of 19 at $N = 70$, none at larger sizes.

The results therefore describe circuits whose MISOC can be stabilized. Both circuits come from the same filtered set of draws, so the paired comparisons are fair, but at small $N$ that set is a minority of random draws.

### D. Letters and re-timing

Each letter is a polyline of 100 stored points, shifted and scaled so that it starts at $(0, 0)$ and ends at $(1, 0)$. For all 52 letters the horizontal extent is exactly $[0, 1]$. One letter width, the unit of every error in the paper, is thus both the start-to-end distance and the horizontal size of the letter. Heights reach up to 1.03 (median 0.66).

Re-timing with cap $k = 3$ gives each segment a duration $\max(1, d_i/(k\,\tilde{d}))$, where $d_i$ is the segment length and $\tilde{d}$ the median segment length. The cumulative durations are normalized to $[0, 1]$, and the pen position at any time is interpolated linearly along the polyline. Because the total is rescaled to the same 500 ms, slowing the fast strokes makes the rest of the letter somewhat faster than in the stored timing. The target used to score a letter is this re-timed trajectory, sampled on the same grid as the pen.

### E. Trial, targets and the hold penalty

The cue is at $t = -1$ s and the go cue at $t = 0$. The ramp is $R(t) = 1 - e^{-(t+1)/\tau_r}$ before go and $R(0)e^{-t/2\,\mathrm{ms}}$ after; in continuous runs it is set exactly to zero 50 ms after go. The arm moves from $(k, 0)$ to $(k + 1, 0)$ during $0 \leq t \leq 0.5$ s and then stays. The network's target is the letter minus the arm, and the readout is asked to match it at 1151 sample times:

- the letter: every 0.5 ms on $[0, 0.5]$ s (1001 samples);
- the hold: 100 samples on $(0.5, 1]$ s, target zero, so the pen parks at the letter's end;
- the preparation: 50 samples on $[-1, -0.02]$ s, target zero.

The hold-speed penalty appends 99 rows $\lambda\,\tau_s(\mathbf{z}_{k+1} - \mathbf{z}_k)/(t_{k+1} - t_k)$, one per pair of consecutive hold samples, with target zero, $\tau_s = 50$ ms and $\lambda = 2$. The step from the last letter sample to the first hold sample is not penalized. All rows enter learning with equal weight, so the letter makes up 80% of all rows (1001 of 1250).

### F. Readout and its gain

The readout pattern $\mathbf{M}_0$ is drawn from a fixed seed, with entries $\mathcal{N}(0, 1/N_E)$ on the excitatory units and zero on the inhibitory units. It is the same for all draws of a given size, and independent across sizes. The gain $s$ in $\mathbf{M} = s\,\mathbf{M}_0$ is set once per draw, with the SOC, in three steps:

1) Compute the linear least-squares states of Eq. (3) of the main text (evoked-energy penalty, no hold-speed rows) for a, b, c and d.
2) Take each state's spread (standard deviation across units) and average over the four letters.
3) Set $s$ to that mean spread divided by 1.5 Hz. Since the states scale as $1/s$, their mean spread becomes 1.5 Hz.

The MISOC of the draw uses the same $\mathbf{M}$. The calibration uses only the linear map and only a–d, so it never sees the nonlinear network's response to the other 48 letters.

### G. Learning one letter

*Parameterization.* A bias pattern is written as $\mathbf{P} = \mathbf{a} - \mathbf{W}g(\mathbf{a})$, which makes $\mathbf{a}$ the fixed point under the constant drive $\mathbf{P}$. The network never reaches $\mathbf{a}$: the ramp is only at $1 - e^{-2.5} = 0.92$ of its final value at go, and the network lags the ramp. At $N = 200$ (MISOC, letter W) the rates at go correlate with $g(\mathbf{a})$ at 0.997 and have 91% of its norm. $\mathbf{a}$ is thus a convenient coordinate for $\mathbf{P}$, not the state the network holds. This costs nothing, because the learning maps below use the real ramp.

*Linear start.* $\mathbf{\Phi}$ maps $\mathbf{a}$ to all sample rows for the linearized network, driven from rest by $R(t)(\mathbf{I} - \mathbf{W})\mathbf{a}$. It is integrated mode by mode over the whole trial, with exact exponential steps of 0.1 ms. If the eigenvector matrix is ill-conditioned (condition number $> 10^8$), the matrix exponential is used instead. The evoked-energy matrix $\mathbf{Q}$ solves $(\mathbf{W} - \mathbf{I})^\top \mathbf{Q} + \mathbf{Q}(\mathbf{W} - \mathbf{I}) = -2\mathbf{I}$ and is scaled to mean eigenvalue 1. With $\mu' = 0.03\,\mathrm{tr}(\mathbf{\Phi}^\top \mathbf{\Phi})/N$, computed from the sample rows alone, the start is $\mathbf{a}_0 = (\tilde{\mathbf{\Phi}}^\top \tilde{\mathbf{\Phi}} + \mu' \mathbf{Q})^{-1} \tilde{\mathbf{\Phi}}^\top \tilde{\mathbf{z}}$, where $\tilde{\mathbf{\Phi}}$ and $\tilde{\mathbf{z}}$ are $\mathbf{\Phi}$ and $\mathbf{z}$ with the hold-speed rows appended.

*Gauss–Newton.* At the best state so far, the nonlinear network is simulated. The exact Jacobian $\mathbf{J}$ of all rows with respect to $\mathbf{a}$ comes from integrating, alongside it and with the same Euler steps, the variational equation $\tau \dot{\mathbf{X}} = -\mathbf{X} + \mathbf{W}\,\mathrm{diag}(g'(\mathbf{x}))\mathbf{X} + R(t)\,(\mathbf{I} - \mathbf{W}\,\mathrm{diag}(g'(\mathbf{a})))$. With residual $\mathbf{e}$, the direction is $\Delta\mathbf{a} = (\mathbf{J}^\top \mathbf{J} + \mu' \mathbf{Q})^{-1} \mathbf{J}^\top \mathbf{e}$.

A candidate $\mathbf{a} + \eta \Delta\mathbf{a}$ is accepted if it raises the score $R^2$ over all sample and penalty rows. This $R^2$ is computed against the mean of each target coordinate, a mean that includes the zero preparation and hold targets. On acceptance $\eta$ grows by half, up to 1, and a new direction is computed; on rejection $\eta$ halves. The search starts at $\eta = 1$ and stops after 20 candidates or after 6 rejections in a row.

*The spread cap.* If a candidate's spread $\mathrm{std}(\mathbf{a})$ exceeds 10 Hz, the whole candidate is rescaled, $\mathbf{a} \leftarrow 10\,\mathbf{a}/\mathrm{std}(\mathbf{a})$. This shrinks the state toward zero rather than clipping it, and it changes the candidate itself, not only the step. The cap is a safeguard against runaway states and almost never binds: the learned spread reaches it (to within 0.01 Hz) for 5 of the 6500 SOC letters (52 letters, 25 draws, 5 sizes) and for none of the MISOC's. The median learned spread is 3.9–5.1 Hz for the SOC and 3.3–4.0 Hz for the MISOC, depending on size. The size of the preparatory states is thus set by the readout gain (calibrated to the 1.5 Hz spread of Hennequin *et al.*) and by the learning itself, not by the cap.

### H. Scoring

Scores come from a separate continuous simulation (`final.run_sequence`) that starts with 300 ms of rest and then runs the trial. For single letters it agrees with the learning simulation to within $10^{-12}$. The paper's RMS error and $R^2$ are computed over the 1001 letter samples only. On the 100-point grid used in early versions, the RMS error is on average 0.13 percentage point higher.

Three further measures are stored but not used as headline results:

- *End error*: the pen's distance from $(k+1, 0)$ at $t = 0.5$ s.
- *Hold deviation*: the largest such distance during the hold.
- *Preparatory excursion*: the pen's largest distance from $(k, 0)$ during preparation.

At $N = 200$ these average 8%, 7% and 3% of a letter width for the SOC, and 5%, 4% and 3% for the MISOC. At $N = 50$ they are 13%, 12% and 7% (SOC) and 10%, 10% and 9% (MISOC). So the pen lands within about a tenth of a letter width of the letter's end and holds there. Preparation is output-null only approximately; the figures do not show this excursion because the pen is drawn as lifted during preparation. Using the model as a continuous pen would require accounting for it.

*Return to rest.* After one letter the input is switched off and the circuit is simulated for 40 s. A circuit counts as settled if, 40 s after the go cue, its mean |rate deviation| over units, averaged over the letters a, l, e and x, is below 0.1 Hz.

### I. Doubled letters and words

Doubled letters and words are each one continuous run, with go cues 4 s apart (1 s preparation, 0.5 s letter, 0.5 s hold and 2 s of rest). Each letter's ramp is the same as in learning. The preparation of letter $k + 1$ starts after the 2 s of rest that follow the hold of letter $k$, from whatever state the circuit is then in, whereas every pattern was learned from rest. The ten words were drawn once from a fixed seed (2026): five letters each, chosen uniformly and with replacement from all 52 letters. A word's score is the mean of its letters' scores, each taken relative to the letter's own arm position. Draws whose MISOC does not return to rest are left out of these analyses (both circuits of the draw).

### J. Input-channel test

The bias is restricted to $\mathbf{P} = \mathbf{B}\mathbf{c}$, where $\mathbf{B}$ has unit-norm columns.

- *Network-derived channels* are the top $K$ right singular vectors of $\mathbf{L} = \mathbf{\Phi}(\mathbf{I} - \mathbf{W})^{-1}$, the linear map from $\mathbf{P}$ to all 1151 sample rows (hold-speed rows excluded). This map is linearized at rest and uses the readout, but no letter.
- *Random channels* are Gaussian vectors from one fixed seed, shared by every draw and both circuits of a size.

The amplitudes $\mathbf{c}$ are learned with the main method's search, except in four respects:

1) The penalty is ridge: $0.03\,\mathrm{tr}(\mathbf{G})/K$ times the identity, with $\mathbf{G} = \mathbf{L}_B^\top \mathbf{L}_B$ for the start and $\mathbf{G} = \mathbf{J}^\top \mathbf{J}$ for each step, where $\mathbf{L}_B$ is $\mathbf{L}\mathbf{B}$ with the hold-speed rows appended (unlike $\mu'$ of the main method, this penalty is computed from all rows).
2) The cap acts on the spread of the *rates* $g(\mathbf{x})$ at the last sample before go, measured by simulation, whereas the main method caps the coordinate $\mathbf{a}$.
3) The cap also applies to the start.
4) A letter whose RMS error exceeds 30% of a letter width after the search is learned again from $0.5\,\mathbf{c}_0$ and, if it still exceeds 30%, from $0.25\,\mathbf{c}_0$, where $\mathbf{c}_0$ is the least-squares start; the fit with the higher $R^2$ over all rows (the search's own score) is kept (`RESTART_SCALES` in `final.py`). A too-strong start can leave the network in a strongly nonlinear range in which no step improves

on it. This was needed for 710 of the 52 000 letter fits of the main text's input-channel results, and afterwards no letter exceeds 30%.

Because the two caps limit different quantities, spreads are not directly comparable between the full-bias and the channel results.

#### K. Robustness runs (held in reserve)

*Noise.* The learned, noise-free patterns are played back with the rate model's Ornstein–Uhlenbeck noise, and nothing is relearned. The noise has $\tau_\xi = 50\,\text{ms}$ and variance $\sigma_0^2(\tau + \tau_\xi)/\tau_\xi$, so an unconnected unit fluctuates with s.d. $\sigma_0$. We use $\sigma_0 = 0.05$, 0.1 and 0.2 Hz, with 10 noise seeds per letter and 3 per word.

*Own readout per draw.* The readout pattern is drawn from seed 1000+ draw seed, the gain is recalibrated, and all letters are relearned.

#### L. Statistics

SOC and MISOC are compared with two-sided Wilcoxon signed-rank tests on per-draw means, paired by draw (the two circuits of a draw share the initial matrix and the readout), with $n = 25$ at each size ($n = 23$ at $N = 200$ for doubled letters and words, whose draws with a MISOC that does not return to rest are left out). Letters within a circuit are not independent samples, so they are averaged before testing. Each size is tested separately at $p < 0.05$, without correction across the five sizes. The abscissa-matched control (Sec. S1) is paired by draw over ten draws. Spreads across draws are sample standard deviations.

#### M. Choices made during development

We reached the final settings in stages; we list them so readers can judge the degrees of freedom.

- *Targets.* Earlier versions asked for the letter alone, or added the hold, the preparation and extra weight on the letter's end.
- *Sampling and filter.* We compared 100 and 200 letter samples with a 50 ms readout filter, and then, at $N = 100$ on a–d, 2.5 against 0.5 ms sampling, with and without the filter.
- *Re-timing.* We added the re-timing (cap 3) after noticing the fast entry and exit strokes.
- *The $\lambda$ test.* $\lambda$ was then chosen by the rule given in the main text, on letters a–d at $N = 100$. Those letters are also the gain-calibration letters, and the test's ten draws are among the 25 draws of the main results. The test is thus separate from the main results in its letters' role, not in its circuits.
- $g_{II}$. We lowered it from 0.8 to 0.5 after the runaway described in the main text, keeping the first ten draws and rebuilding only their MISOCs.
- *Never tuned on the main results:* the model, the circuit recipe, the regularization (0.03), the iteration limits and the draw selection.
